# A highly contiguous genome assembly for the Yellow Warbler (*Setophaga petechia*)

**DOI:** 10.1101/2024.01.10.575107

**Authors:** Whitney L. E. Tsai, Merly Escalona, Kimball L. Garrett, Ryan S. Terrill, Ruta Sahasrabudhe, Oanh Nguyen, Eric Beraut, William Seligmann, Colin W. Fairbairn, Ryan J. Harrigan, John E. McCormack, Michael E. Alfaro, Thomas B. Smith, Rachael A. Bay

## Abstract

The Yellow Warbler (*Setophaga petechia*) is a small songbird in the New World Warbler family (Parulidae) that exhibits phenotypic and ecological differences across a widespread distribution and is important to California’s riparian habitat conservation. Here, we present a high-quality *de novo* genome assembly of a vouchered female Yellow Warbler from southern California. Using HiFi long-read and Omni-C proximity sequencing technologies, we generated a 1.22 Gb assembly including 687 scaffolds with a contig N50 of 6.80 Mb, scaffold N50 of 21.18 Mb, and a BUSCO completeness score of 96.0%. This highly contiguous genome assembly provides an essential resource for understanding the history of gene flow, divergence, and local adaptation and can inform conservation management of this charismatic bird species.

## Introduction

The Yellow Warbler (*Setophaga petechia*) is a widespread songbird species distributed from Alaska to northern South America (Figure 1). The species complex comprises up to 43 subspecies in four distinct subspecies groups that display notable diversity in phenotype and ecology across their range (Browning, 1994; Klein & Brown, 1994; Salgado-Ortiz et al., 2008; Wilson & Holberton, 2004). This phenotypic diversity and the presence of both migratory and resident populations have encouraged investigation into the history of adaptation, divergence, and gene flow in this species (Chavarria-Pizarro et al., 2019; Chaves et al., 2012; Gibbs et al., 2000; Machkour-M’Rabet et al., 2023; Milot et al., 2000). Additionally, as a widespread migratory bird species, the Yellow Warbler inhabits variable environmental conditions across its range, allowing for the investigation into the influence of climate on geographic variation and genomic capacity to adapt to climate change (Bay et al., 2018; Chen et al., 2022; DeSaix et al., 2022).

**Figure 1.**
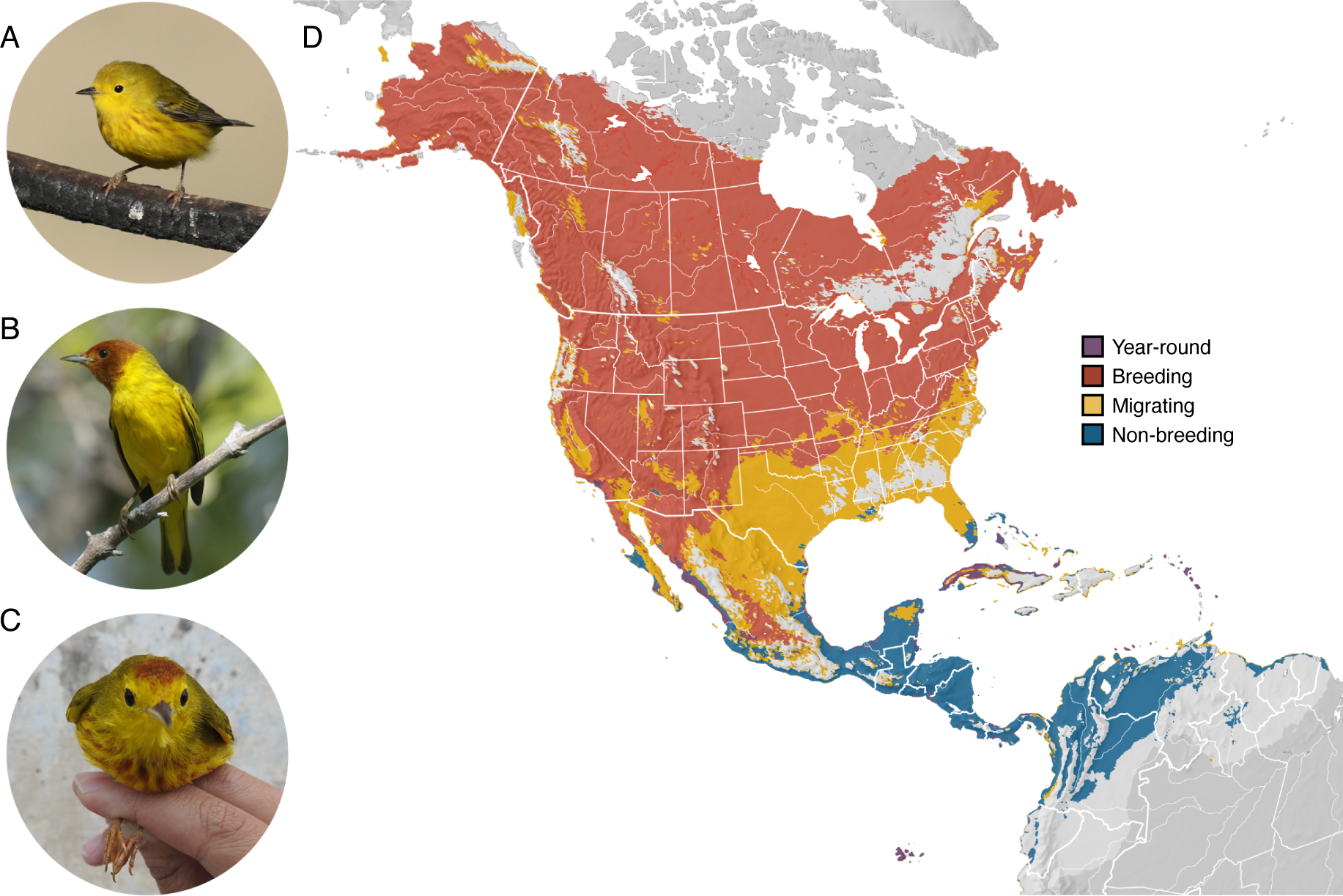
Geographic variation and distribution of Yellow Warblers (*Setophaga petechia*). A) The Northern (*aestiva*) group includes migratory subspecies with chestnut streaking on the breast. Northern subspecies breed in North America and winter in Central and northern South America. Photo taken by R. S. Terrill at Piute Ponds, Los Angeles, CA, USA. B) The Mangrove (*erithachorides*) group includes resident subspecies with a characteristic all chestnut head. Mangrove subspecies inhabit mangroves along the coasts of Central and northern South America year-round. Photo taken by R. S. Terrill on Isla Holbox, Quintana Roo, MX. C) The Galapagos (*aureola*) and Golden (*petechia*) subspecies groups includes resident subspecies with a chestnut cap and thick breast streaking except for *S. p. ruficapilla* from Martinique which exhibits the Mangrove phenotype. Populations of the Galapagos subspecies are found on the Galapagos Islands and Cocos Island off Costa Rica and Golden subspecies are found on the islands of the Caribbean. Photo taken by W. L. E. Tsai on Isla Cozumel, Quintana Roo, MX. D) Map of species distributional abundance (Fink et al., 2022). Shaded colors indicate seasonal shifts in distributions: year-round (purple), breeding (red), migrating (yellow), and non-breeding (blue).

In California, Yellow Warblers are listed as a Species of Special Concern (Shuford et al., 2008) and have experienced notable declines over the last 50 years (Sauer et al., 2014). Previous genomic work indicates that the inability to adapt to climate change may play a role in population declines in California (Bay et al., 2018). California wetlands and riparian corridors are crucial stopover and breeding habitats for Yellow Warblers and other species of migratory birds. In the last century, 90-95% of historic wetland and riparian habitats have been lost, and those that remain are threatened by development and climate change (Dahl, 1990; Krueper, 1996; Poff et al., 2012). As indicators of healthy riparian habitat, understanding how California Yellow Warbler populations adapt to dramatic changes in their environment will inform conservation action and help mitigate habitat loss in other vulnerable and threatened riparian species, like the California Red-legged Frog (*Rana draytonii*), the Riparian Brush Rabbit (*Sylvilagus bachmani riparius)*, and the Valley Elderberry Longhorn Beetle (*Desmocerus californicus dimorphus)* (Collinge et al., 2001; Davidson et al., 2001; Heath & Ballard, 2003; Phillips et al., 2005).

The evolutionary and conservation genomics studies needed to address these questions increasingly rely on low-coverage, whole genome sequencing (WGS), which requires a high-quality reference genome for alignment. Reference genome assemblies provide a map of the structural features and organization of the genome and the choice of reference genome assembly for WGS studies can impact evolutionary inferences like demographic history and genetic diversity (Gopalakrishnan et al., 2017). Currently, there are four genome assemblies generated with short-read sequencing technology for the genus *Setophaga*. There is one Yellow-rumped Warbler (*S. coronata*) chromosome-level assembly (Toews et al., 2016), two Kirtland’s Warbler (*S. kirtlandii*) scaffold-level assemblies (Feng et al., 2020), and the existing draft genome assembly for Yellow Warbler has a length of 1.26 Gb, a total of 18,414 scaffolds, and a scaffold N50 491.7 kb (Bay et al., 2018). The use of an interspecific reference genome assembly can lead to many errors and biases, including lower mapping ability (especially in regions with higher evolutionary rates) and inaccurate gene order (Prasad et al., 2022). The high number and relatively short scaffold length of the existing Yellow Warbler genome assembly could hinder the identification of structural variants often maintained between and within species and are important in adaptive evolution, speciation, and generating morphological diversity (Lamichhaney et al., 2016; Mérot et al., 2020; Wellenreuther & Bernatchez, 2018). Additionally, reference genome assemblies generated solely from short-read sequencing technology fail to resolve lengths and placement of repeat regions, such as transposable elements or telomeres, leading to gaps in avian genome assemblies (Peona et al., 2021). This highlights the need for a high-quality, species-specific reference genome for WGS studies.

Here, we present a new genome assembly for the Yellow Warbler generated as part of the California Conservation Genomics Project (CCGP) consortium (Shaffer et al., 2022). We used high-molecular-weight (HMW) genomic DNA (gDNA) extracted from a vouchered, female bird collected in California and leveraged Pacific Biosciences (PacBio) HiFi long-read and Dovetail Genomics Omni-C proximity sequencing technologies. This produced a high-quality genome assembly that will allow us to better understand evolutionary processes like phenotypic variation and migration and conduct conservation genomics studies to inform conservation initiatives.

## Methods

### Biological Materials

We sampled muscle, liver, and other tissues from a female Yellow Warbler collected using mist nets near Stephen Sorensen Park (34.60549°N, 117.8306°W) in Los Angeles County, California on September 25, 2020. This migrant Yellow Warbler can presumably be assigned to *S. p. brewsteri* based on collection date and locality (Browning, 1994) and was collected with approval from the following entities: California Department of Fish and Wildlife Scientific Collecting Permit (#SC-000939), US Fish and Wildlife Services Scientific Collecting Permit (MB708062-0), and US Geological Survey Banding Permit (22804-B). Tissue samples were retrieved and flash-frozen in liquid nitrogen, and the first muscle tissues were frozen within two minutes of specimen collection. A voucher specimen and tissue are deposited at the Natural History Museum of Los Angeles (LACM Bird #122168, KLG4550, LAF9440). Additional tissues for this individual are housed in the CCGP tissue repository at the University of California, Los Angeles under identification YEWA_CCGP3.

### Nucleic acid extraction, library preparation, and sequencing

We extracted HMW gDNA from 30mg of flash-frozen heart tissue. We homogenized the tissue by grinding it in a mortar and pestle in liquid nitrogen. We lysed the homogenized tissue at room temperature overnight with 2 ml of lysis buffer containing 100mM NaCl, 10 mM Tris-HCl pH 8.0, 25 mM EDTA, 0.5% (w/v) SDS, and 100µg/ml Proteinase K. We treated the lysate with 20µg/ml RNAse at 37^0^C for 30 minutes. We cleaned the lysate with equal volumes of phenol/chloroform using phase lock gels (Quantabio, MA; Cat # 2302830). We precipitated the DNA from the cleaned lysate by adding 0.4X volume of 5M ammonium acetate and 3X volume of ice-cold ethanol. We washed the pellet twice with 70% ethanol and resuspended it in elution buffer (10mM Tris, pH 8.0). We measured DNA purity using absorbance ratios (260/280 = 1.87 and 260/230 = 2.29) using a NanoDrop ND-1000 spectrophotometer. We quantified DNA yield (30µg) using a Qubit 2.0 Fluorometer (Thermo Fisher Scientific, MA). We verified HMW gDNA integrity on a Femto pulse system (Agilent Technologies, CA), where 80% of the DNA was found in fragments above 120 Kb.

According to the manufacturer’s instructions, we constructed the HiFi Single Molecule, Real-Time (SMRT) library using SMRTbell Express Template Prep Kit v2.0 (PacBio, CA; Cat. #100-938-900). We sheared HMW gDNA to a target DNA size distribution between 15 – 20 kb and concentrated it using 0.45X of AMPure PB beads (PacBio; Cat. #100-265-900). We performed the enzymatic incubations as follows: removal of single-strand overhangs at 37°C for 15 minutes, DNA damage repair at 37°C for 30 minutes, end repair at 20°C for 10 minutes, A-tailing at 65°C for 30 minutes, ligation of overhang adapter v3 at 20°C for 60 minutes, ligase inactivation at 65°C for 10 minutes, and nuclease treatment at 37°C for 1 hour. We purified and concentrated the library with 0.45X Ampure PB beads for size selection to collect fragments greater than 7-9 kb using the BluePippin/PippinHT system (Sage Science, MA; Cat #BLF7510/HPE7510). The HiFi library averaged 15 – 20 kb. It was sequenced at UC Davis DNA Technologies Core (Davis, CA) using two 8M SMRT cells, Sequel II sequencing chemistry 2.0, and 30-hour movies each on a PacBio Sequel II sequencer.

We used the Omni-C^TM^ Kit (Dovetail Genomics, CA) for Omni-C proximity sequencing according to the manufacturer’s protocol with slight modifications. First, we ground muscle tissue (Sample YEWA_CCGP3; LACM Bird #122168, KLG4550, LAF9440) with a mortar and pestle while cooled with liquid nitrogen. Subsequently, chromatin was fixed in place in the nucleus. We passed the suspended chromatin solution through 100 μm and 40 μm cell strainers to remove large debris. We digested fixed chromatin under various conditions of DNase I until a suitable fragment length distribution of DNA molecules was obtained. We repaired chromatin ends, ligated a biotinylated bridge adapter, and performed proximity ligation of adapter-containing ends. After proximity ligation, crosslinks were reversed, and the DNA was purified from proteins. We treated purified DNA to remove biotin that was not internal to ligated fragments. We generated a next-generation sequencing library using an NEB Ultra II DNA Library Prep kit (New England Biolabs, MA) with an Illumina-compatible y-adapter. Then, we captured biotin-containing fragments using streptavidin beads. We split the post-capture product into two replicates before PCR enrichment to preserve library complexity, with each replicate receiving unique dual indices. The library was sequenced at the Vincent J. Coates Genomics Sequencing Lab (Berkeley, CA) on an Illumina NovaSeq platform 6000 (Illumina, CA) to generate approximately 100 million 2 x 150 bp read pairs per GB genome size.

### Nuclear genome assembly

We assembled the Yellow Warbler genome following the CCGP assembly pipeline Version 4.0, as outlined in Table 1, which lists the tools and non-default parameters used in the assembly. The pipeline uses PacBio HiFi reads and Omni-C data to produce high-quality and highly contiguous genome assemblies, minimizing manual curation. We removed remnant adapter sequences from the PacBio HiFi dataset using HiFiAdapterFilt (Sim et al., 2022). Then, we obtained the initial phased diploid assembly using HiFiasm (Cheng et al., 2022) with the filtered PacBio HiFi reads and the Omni-C dataset. We aligned the Omni-C data to both assemblies following the Arima Genomics Mapping Pipeline (https://github.com/ArimaGenomics/mapping_pipeline) and then scaffolded both assemblies with SALSA (Ghurye et al., 2017, 2019).

**Table 1.**
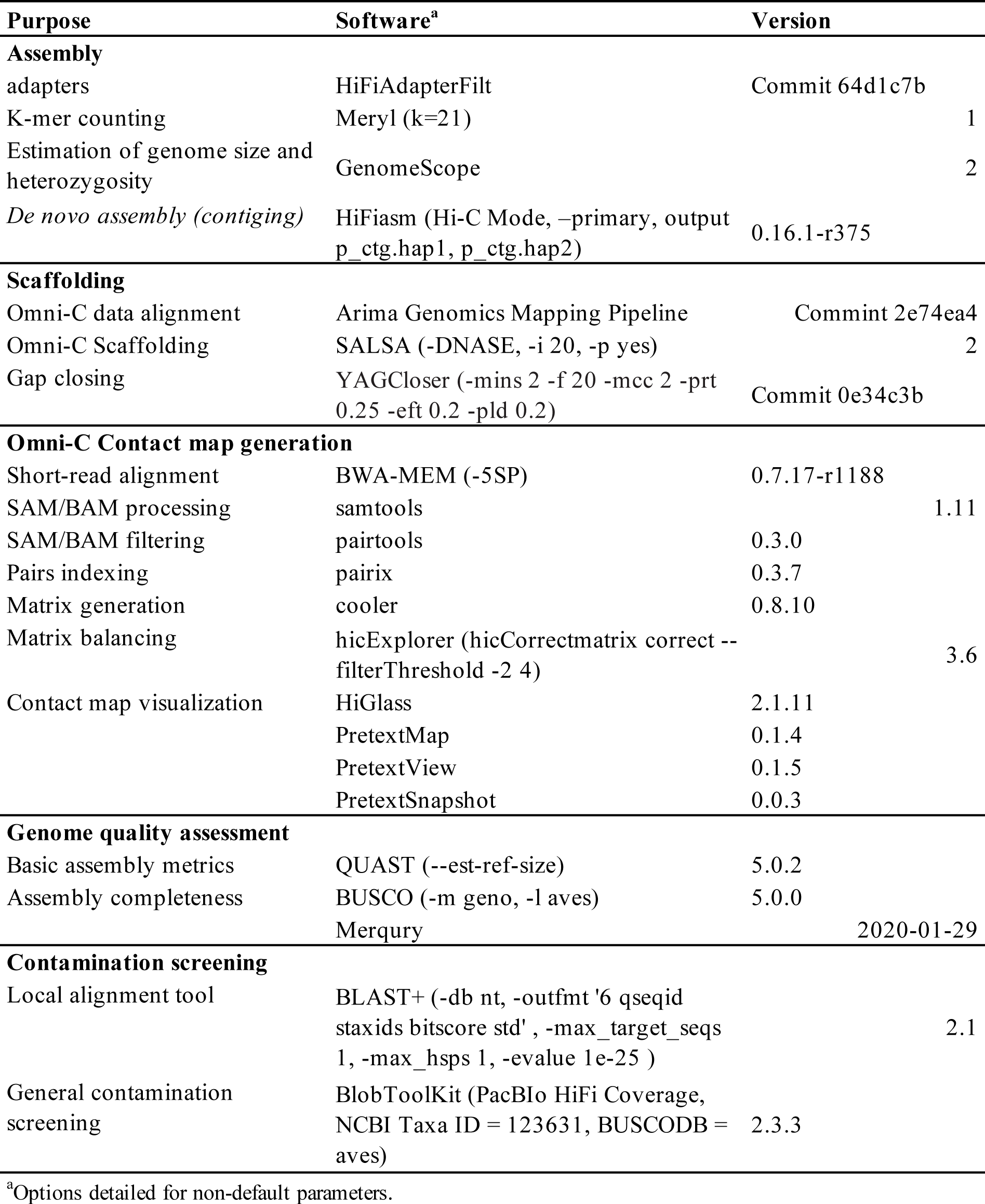
Assembly pipeline and software used for assembly of the Yellow Warbler genome. Software citations are listed in the text.

We generated Omni-C contact maps for both assemblies by aligning the Omni-C data with BWA-MEM (Li, 2013), identified ligation junctions, and generated Omni-C pairs using pairtools (Goloborodko et al., 2018). We generated a multi-resolution Omni-C matrix with a cooler (Abdennur & Mirny, 2020) and balanced it with hicExplorer (Ramírez et al., 2018). We used HiGlass [Version 2.1.11] (Kerpedjiev et al., 2018) and the PretextSuite (https://github.com/wtsi-hpag/PretextView; https://github.com/wtsi-hpag/PretextMap; https://github.com/wtsi-hpag/PretextSnapshot) to visualize the contact maps and then we checked the contact maps for major mis-assemblies. In detail, if we identified a strong off-diagonal signal and a lack of signal in the consecutive genomic region in the proximity of a join made by the scaffolder, we dissolved it by breaking the scaffolds at the coordinates of the join. After this process, no further manual joins were made. Some remaining gaps (joins generated by the scaffolder) were closed using the PacBio HiFi reads and YAGCloser (https://github.com/merlyescalona/yagcloser). Finally, we checked for contamination using the BlobToolKit Framework (Challis et al., 2020). Given the fragmentation of the assemblies, these were tagged as primary or alternate based on overall metrics.

### Genome assembly assessment

We generated k-mer counts from the PacBio HiFi reads using meryl (https://github.com/marbl/meryl). The k-mer database was then used in GenomeScope2.0 (Ranallo-Benavidez et al., 2020) to estimate genome features, including genome size, heterozygosity, and repeat content. To obtain general contiguity metrics, we ran QUAST (Gurevich et al., 2013). To evaluate genome quality and functional completeness, we used BUSCO (Manni et al., 2021) with the Aves ortholog database (aves_odb10) containing 8,338 genes. Base level accuracy (QV) and k-mer completeness were assessed using the previously generated meryl database and merqury (Rhie et al., 2020). We further estimated genome assembly accuracy via BUSCO gene set frameshift analysis using the pipeline described in (Korlach et al., 2017). Measurements of the size of the phased blocks are based on the size of the contigs generated by HiFiasm on HiC mode. We follow the quality metric nomenclature established by (Rhie et al., 2021), with the genome quality code x.y. P.Q.C, where x = log10[contig NG50]; y = log10[scaffold NG50]; P = log10 [phased block NG50]; Q = Phred base accuracy QV (quality value); C = % genome represented by the first ‘n’ scaffolds, following a known karyotype of 2n =80 for *S. petechia* (Bird Chromosome Database, Chromosome number data V3.0/2022 - (Degrandi et al., 2020; Hobart, 1991)). Quality metrics for the notation were calculated on the primary assembly (bSetPet1.0.p).

### Mitochondrial genome assembly

We assembled the mitochondrial genome of *S. petechia* from the PacBio HiFi reads using the reference-guided pipeline MitoHiFi (Allio et al., 2020; Uliano-Silva et al., 2021). We used the mitochondrial sequence of *Setophaga kirtlandii* (NCBI:NC_051027.1) as the starting reference sequence. After completion of the nuclear genome, we searched for matches of the resulting mitochondrial assembly sequence in the nuclear genome assembly using BLAST+ (Camacho et al., 2009) and filtered out contigs and scaffolds from the nuclear genome with a percentage of sequence identity >99% and size smaller than the mitochondrial assembly sequence.

## Results

### Sequencing Data

The Omni-C and PacBio HiFi sequencing libraries generated 85.3 million read pairs and 2.7 million reads, respectively. The latter yielded 40.87-fold coverage (N50 read length 17,523 bp; minimum read length 41 bp; mean read length 17,110 bp; maximum read length of 54,497 bp). Based on PacBio HiFi reads, we estimated a genome assembly size of 1.14 Gb, 0.245% sequencing error rate and 1.16% nucleotide heterozygosity rate using Genomescope2.0. The k-mer spectrum based on PacBio HiFi reads shows (Figure 2A) a bimodal distribution with two major peaks at 19- and 39-fold coverage, where peaks correspond to homozygous and heterozygous states of a diploid species.

**Figure 2.**
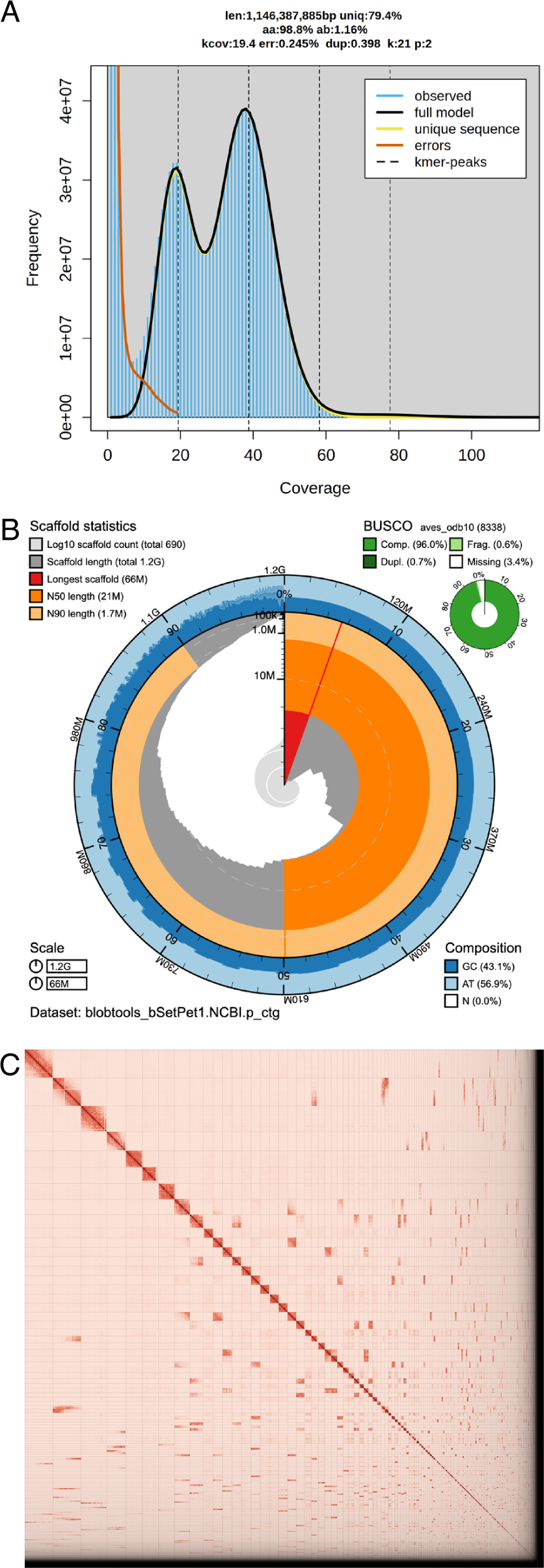
Visual overview of genome assembly metrics. A) Kmer spectra output generated from PacBio HiFi data without adapters using GenomScope2.0. The bimodal pattern observed corresponds to a diploid genome. K-mers covered at lower coverage and lower frequency correspond to differences between haplotypes, whereas the higher coverage and higher frequency k-mers correspond to the similarities between haplotypes. B) BlobToolKit Snail plot showing a graphical representation of the quality metrics presented in Table 2 for the *Setophaga petechia* primary assembly (bSetPet1.0.p). The plot circle represents the full size of the assembly. From the inside-out, the central plot covers scaffold and length-related metrics. The central light gray spiral shows the cumulative scaffold count with a white line at each order of magnitude. The red line represents the size of the longest scaffold; all other scaffolds are arranged in size-order moving clockwise around the plot and drawn in gray starting from the outside of the central plot. Dark and light orange arcs show the scaffold N50 and scaffold N90 values. The outer light and dark blue ring show the mean, maximum, and minimum GC versus AT content at 0.1% intervals (Challis et al. 2020). C) Omni-C contact map for the primary genome assembly generated with PretextSnapshot. Omni-C contact maps translate proximity of genomic regions in 3D space to contiguous linear organization. Each cell in the contact map corresponds to sequencing data supporting the linkage (or join) between 2 such regions. Scaffolds are separated by black lines and higher density corresponds to higher levels of fragmentation.

**Table 2.**
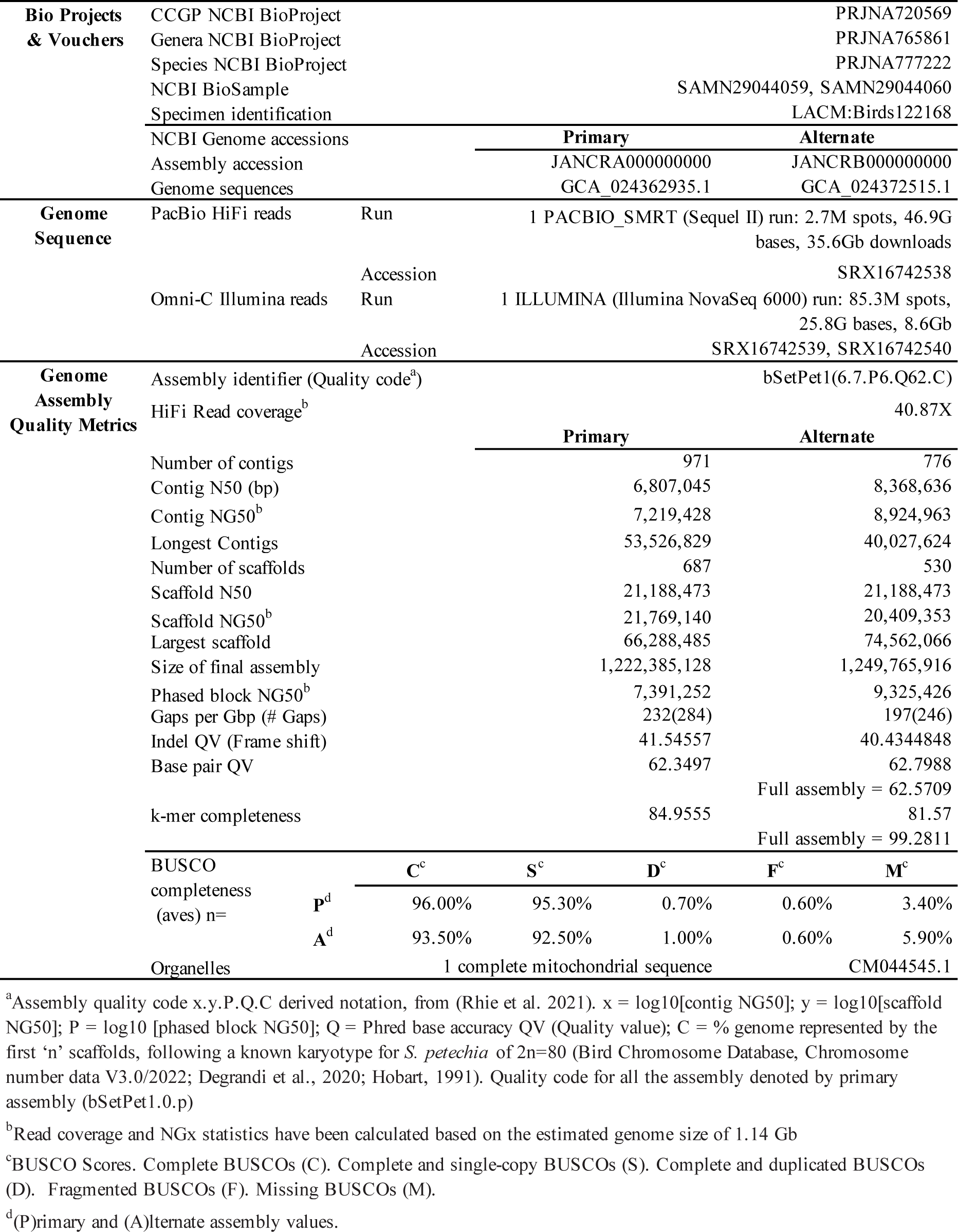
Sequencing and assembly statistics and accession information for the primary and alternate assemblies of the Yellow Warbler (*Setophaga petechia*) genome.

### Nuclear genome assembly

The final assembly consists of two haplotypes tagged as primary and alternate (bSetPet1.0.p and bSetPet1.0.a). Both genome assembly sizes are similar but not equal to the estimated value from Genomescope2.0 (Figure 2A). The primary assembly (bSetPet1.0.p) consists of 687 scaffolds spanning 1.22 Gb with contig N50 of 6.8 Mb, scaffold N50 of 21.18 Mb, longest contig of 53.52 Mb, and largest scaffold of 66.28 Mb. The alternate assembly (bSetPet1.0.a) consists of 530 scaffolds, spanning 1.24 Gb with contig N50 of 8.3Mb, scaffold N50 of 21.18 Mb, largest contig 40.02 Mb and largest scaffold of 74.56 Mb. The Omni-C contact maps suggest highly contiguous primary and alternate assemblies (Figure 2C and Supplementary Figure S1B). The primary assembly has a BUSCO completeness score of 96.0% using the Aves gene set, a per-base quality (QV) of 62.34, a k-mer completeness of 84.95, and a frameshift indel QV of 41.54. In comparison, the alternate assembly has a BUSCO completeness score of 93.5% using the same gene set, a per-base quality (QV) of 62.79, a k-mer completeness of 81.57, and a frameshift indel QV of 40.43.

During manual curation, we identified 13 misassemblies requiring breaking nine joins on the primary assembly and four on the alternate assembly. We were able to close a total of five gaps, three on the primary and two on the alternate assembly. We removed two contigs, one per assembly, corresponding to mitochondrial contaminants. Detailed assembly statistics are reported in Table 2, and a graphical representation of the primary assembly in Figure 2B (see Supplementary Figure S1A for the alternate assembly). We have deposited both assemblies on NCBI (See Table 2 and Data Availability for details).

### Mitochondrial genome assembly

We assembled a mitochondrial genome with MitoHiFi. The final mitochondrial assembly has a size of 16,809 bp. The base composition of the final assembly version is A=30.19%, C=31.77%, G=14.19%, T=23.85%, and consists of 22 unique transfer RNAs and 13 protein-coding genes.

## Discussion

Here, we present a highly contiguous genome assembly for the Yellow Warbler with two pseudo haplotypes. Our genome assemblies meet thresholds for proposed quality standards for vertebrate and avian genomes (Jarvis, 2016; Kapusta & Suh, 2017; Rhie et al., 2021). Compared to the existing *Setophaga* genomes, the primary Yellow Warbler genome assembly presented here has the highest BUSCO completeness (96.0% of Aves orthologs present) and the highest contig N50 (6.8 Mb). Although the Yellow-rumped and Kirtland’s Warbler genome assemblies have higher scaffold N50 values, our Yellow Warbler genome assembly has the fewest gaps greater than 5 N’s (284 compared to 49-67K in other *Setophaga* genome assemblies), which highlights the improvement gained when using long-read sequencing technology in combination with short reads for more contiguous and complete genomes.

The reference genome presented here provides an essential resource for evolutionary research and conservation efforts in California and beyond. Future range-wide genomic analyses will facilitate investigations into the history of gene flow and divergence between the various subspecies groups in this complex (Browning, 1994; Chaves et al., 2012; Machkour-M’Rabet et al., 2023). This system-wide genomic context lends itself to investigations into the genetic basis underlying both phenotypic diversity and the evolution of migration (Aguillon et al., 2021; Caballero-López et al., 2022; Delmore et al., 2020; Franchini et al., 2017; Toews et al., 2016). Future landscape genomic analyses investigating environmental associations with genomic variation could identify loci important for local adaptation in this widespread species (Bay et al., 2018; Chen et al., 2022; Forester et al., 2018). Using this framework with future climate models will allow for predictions of how Yellow Warblers may adapt to future climate change and identify both populations that are likely to persist in and vulnerable to future climate change regimes, which will guide local conservation implementation (Fitzpatrick & Keller, 2015; Shaffer et al., 2022). This will be especially important for California populations experiencing population declines and dwindling breeding habitat, which could benefit from direct conservation and management efforts (Heath & Ballard, 2003; Shuford et al., 2008). Overall, the Yellow Warbler genome presented here provides a key resource for investigating phenotypic and ecological evolution and conservation in this charismatic migratory bird species.

## Acknowledgements

PacBio Sequel II library prep and sequencing was carried out at the DNA Technologies and Expression Analysis Cores at the UC Davis Genome Center, supported by NIH Shared Instrumentation Grant 1S10OD010786-01. Deep sequencing of Omni-C libraries used the Novaseq S4 sequencing platforms at the Vincent J. Coates Genomics Sequencing Laboratory at UC Berkeley, supported by NIH S10 OD018174 Instrumentation Grant. We thank the staff at the UC Davis DNA Technologies and Expression Analysis Cores and the UC Santa Cruz Paleogenomics Laboratory for their diligence and dedication to generating high quality sequence data. We thank Maeve Secor for help with fieldwork; Tara Luckau, Dr. Courtney Miller, and Dr. Erin Toffelmier for help with coordination and sample submission.

## Data Availability

Data generated for this study are available under NCBI BioProject PRJNA777222. Raw sequencing data for individual with voucher LACM:122168 (NCBI BioSamples SAMN29044059, SAMN29044060) are deposited in the NCBI Short Read Archive (SRA) under SRX16742538 for PacBio HiFi sequencing data, and SRX16742539 and SRX16742540 for the Omni-C Illumina sequencing data. GenBank accessions for both primary and alternate assemblies are GCA_024362935.1 and GCA_024372515.1; and for genome sequences JANCRA000000000 and JANCRB000000000. The GenBank organelle genome assembly for the mitochondrial genome is CM044545.1. Assembly scripts and other data for the analyses presented can be found at the following GitHub repository: www.github.com/ccgproject/ccgp_assembly

## Supplementary Material

**Figure S1.** Visual overview of genome assembly metrics for alternate assembly (bSetPet1.0.a).

## References

Abdennur, N., & Mirny, L. A. (2020). Cooler: Scalable storage for Hi-C data and other genomically labeled arrays. Bioinformatics, 36(1), 311–316. 10.1093/bioinformatics/btz540

Aguillon, S. M., Walsh, J., & Lovette, I. J. (2021). Extensive hybridization reveals multiple coloration genes underlying a complex plumage phenotype. Proceedings of the Royal Society B: Biological Sciences, 288(1943), 20201805. 10.1098/rspb.2020.1805

Allio, R., Schomaker-Bastos, A., Romiguier, J., Prosdocimi, F., Nabholz, B., & Delsuc, F. (2020). MitoFinder: Efficient automated large-scale extraction of mitogenomic data in target enrichment phylogenomics. Molecular Ecology Resources, 20(4), 892–905. 10.1111/1755-0998.13160

Bay, R. A., Harrigan, R. J., Underwood, V. L., Gibbs, H. L., Smith, T. B., & Ruegg, K. (2018). Genomic signals of selection predict climate-driven population declines in a migratory bird. Science, 359(6371), 83–86. 10.1126/science.aan4380

Browning, M. R. (1994). A taxonomic review of Dendroica petechia (Yellow Warbler) (Aves: Parulinae). PROCEEDINGS OF THE BIOLOGICAL SOCIETY OF WASHINGTON, 107(1), 27–51.

Caballero-López, V., Lundberg, M., Sokolovskis, K., & Bensch, S. (2022). Transposable elements mark a repeat-rich region associated with migratory phenotypes of willow warblers (Phylloscopus trochilus). Molecular Ecology, 31(4), 1128–1141. 10.1111/mec.16292

Camacho, C., Coulouris, G., Avagyan, V., Ma, N., Papadopoulos, J., Bealer, K., & Madden, T. L. (2009). BLAST+: Architecture and applications. BMC Bioinformatics, 10(1), 421. 10.1186/1471-2105-10-421

Challis, R., Richards, E., Rajan, J., Cochrane, G., & Blaxter, M. (2020). BlobToolKit – Interactive Quality Assessment of Genome Assemblies. G3 Genes|Genomes|Genetics, 10(4), 1361–1374. 10.1534/g3.119.400908

Chavarria-Pizarro, T., Gomez, J. P., Ungvari-Martin, J., Bay, R., Miyamoto, M. M., & Kimball, R. (2019). Strong phenotypic divergence in spite of low genetic structure in the endemic Mangrove Warbler subspecies (Setophaga petechia xanthotera) of Costa Rica. Ecology and Evolution, n/a(n/a). 10.1002/ece3.5826

Chaves, J. A., Parker, P. G., & Smith, T. B. (2012). Origin and population history of a recent colonizer, the yellow warbler in Galápagos and Cocos Islands. Journal of Evolutionary Biology, 25(3), 509–521. 10.1111/j.1420-9101.2011.02447.x

Chen, Y., Jiang, Z., Fan, P., Ericson, P. G. P., Song, G., Luo, X., Lei, F., & Qu, Y. (2022). The combination of genomic offset and niche modelling provides insights into climate change-driven vulnerability. Nature Communications, 13(1), Article 1. 10.1038/s41467-022-32546-z

Cheng, H., Jarvis, E. D., Fedrigo, O., Koepfli, K.-P., Urban, L., Gemmell, N. J., & Li, H. (2022). Haplotype-resolved assembly of diploid genomes without parental data. Nature Biotechnology, 40(9), 1332–1335. 10.1038/s41587-022-01261-x

Collinge, S. K., Holyoak, M., Barr, C. B., & Marty, J. T. (2001). Riparian habitat fragmentation and population persistence of the threatened valley elderberry longhorn beetle in central California. Biological Conservation, 100(1), 103–113. 10.1016/S0006-3207(00)00211-1

Dahl, T. E. (1990). Wetlands losses in the United States, 1780’s to 1980’s. Report to the Congress (PB-91-169284/XAB). National Wetlands Inventory, St. Petersburg, FL (USA). https://www.osti.gov/biblio/5527872-wetlands-losses-united-states-report-congress

Davidson, C., Bradley Shaffer, H., & Jennings, M. R. (2001). Declines of the California Red-Legged Frog: Climate, Uv-B, Habitat, and Pesticides Hypotheses. Ecological Applications, 11(2), 464–479. 10.1890/1051-0761(2001)011[0464:DOTCRL]2.0.CO;2

Degrandi, T. M., Barcellos, S. A., Costa, A. L., Garnero, A. D. V., Hass, I., & Gunski, R. J. (2020). Introducing the Bird Chromosome Database: An Overview of Cytogenetic Studies in Birds. Cytogenetic and Genome Research, 160(4), 199–205. 10.1159/000507768

Delmore, K., Illera, J. C., Pérez-Tris, J., Segelbacher, G., Lugo Ramos, J. S., Durieux, G., Ishigohoka, J., & Liedvogel, M. (2020). The evolutionary history and genomics of European blackcap migration. eLife, 9, e54462. 10.7554/eLife.54462

DeSaix, M. G., George, T. L., Seglund, A. E., Spellman, G. M., Zavaleta, E. S., & Ruegg, K. C. (2022). Forecasting climate change response in an alpine specialist songbird reveals the importance of considering novel climate. Diversity and Distributions, 28(10), 2239– 2254. 10.1111/ddi.13628

Feng, S., Stiller, J., Deng, Y., Armstrong, J., Fang, Q., Reeve, A. H., Xie, D., Chen, G., Guo, C., Faircloth, B. C., Petersen, B., Wang, Z., Zhou, Q., Diekhans, M., Chen, W., Andreu-Sánchez, S., Margaryan, A., Howard, J. T., Parent, C., … Zhang, G. (2020). Dense sampling of bird diversity increases power of comparative genomics. Nature, 587(7833), Article 7833. 10.1038/s41586-020-2873-9

Fink, D., Auer, T., Johnston, A., Strimas-Mackey, M., Ligocki, S., Robinson, O., Hochachka, W., Jaromczyk, L., Rodewald, A., Wood, C., Davies, I., & Spencer, A. (2022). eBird Status and Trends, Data Version: 2021; Released: 2022. Cornell Lab of Ornithology, Ithaca, New York. 10.2173/ebirdst.2021

Fitzpatrick, M. C., & Keller, S. R. (2015). Ecological genomics meets community-level modelling of biodiversity: Mapping the genomic landscape of current and future environmental adaptation. Ecology Letters, 18(1), 1–16. 10.1111/ele.12376

Forester, B. R., Lasky, J. R., Wagner, H. H., & Urban, D. L. (2018). Comparing methods for detecting multilocus adaptation with multivariate genotype–environment associations. Molecular Ecology, 27(9), 2215–2233. 10.1111/mec.14584

Franchini, P., Irisarri, I., Fudickar, A., Schmidt, A., Meyer, A., Wikelski, M., & Partecke, J. (2017). Animal tracking meets migration genomics: Transcriptomic analysis of a partially migratory bird species. Molecular Ecology, 26(12), 3204–3216. 10.1111/mec.14108

Ghurye, J., Pop, M., Koren, S., Bickhart, D., & Chin, C.-S. (2017). Scaffolding of long read assemblies using long range contact information. BMC Genomics, 18(1), 527. 10.1186/s12864-017-3879-z

Ghurye, J., Rhie, A., Walenz, B. P., Schmitt, A., Selvaraj, S., Pop, M., Phillippy, A. M., & Koren, S. (2019). Integrating Hi-C links with assembly graphs for chromosome-scale assembly. PLOS Computational Biology, 15(8), e1007273. 10.1371/journal.pcbi.1007273

Gibbs, H. L., Dawson, R. J. G., & Hobson, K. A. (2000). Limited differentiation in microsatellite DNA variation among northern populations of the yellow warbler: Evidence for male-biased gene flow? Molecular Ecology, 9(12), 2137–2147. 10.1046/j.1365-294X.2000.01136.x

Goloborodko, A., Abdennur, N., Venev, S., hbbrandao, & gfudenberg. (2018). mirnylab/pairtools: V0.2.0 [Computer software]. Zenodo. 10.5281/zenodo.1490831

Gopalakrishnan, S., Samaniego Castruita, J. A., Sinding, M.-H. S., Kuderna, L. F. K., Räikkönen, J., Petersen, B., Sicheritz-Ponten, T., Larson, G., Orlando, L., Marques-Bonet, T., Hansen, A. J., Dalén, L., & Gilbert, M. T. P. (2017). The wolf reference genome sequence (Canis lupus lupus) and its implications for Canis spp. Population genomics. BMC Genomics, 18(1), 495. 10.1186/s12864-017-3883-3

Gurevich, A., Saveliev, V., Vyahhi, N., & Tesler, G. (2013). QUAST: Quality assessment tool for genome assemblies. Bioinformatics, 29(8), 1072–1075. 10.1093/bioinformatics/btt086

Heath, S. K., & Ballard, G. (2003). Patterns of Breeding Songbird Diversity and Occurrence in Riparian Habitats of the Eastern Sierra Nevada.

Hobart, H. H. (1991). Comparative karyology in nine-primaried oscines (Aves). https://repository.arizona.edu/handle/10150/185492

Jarvis, E. D. (2016). Perspectives from the Avian Phylogenomics Project: Questions that Can Be Answered with Sequencing All Genomes of a Vertebrate Class. Annual Review of Animal Biosciences, 4(1), 45–59. 10.1146/annurev-animal-021815-111216

Kapusta, A., & Suh, A. (2017). Evolution of bird genomes—A transposon’s-eye view. Annals of the New York Academy of Sciences, 1389(1), 164–185. 10.1111/nyas.13295

Kerpedjiev, P., Abdennur, N., Lekschas, F., McCallum, C., Dinkla, K., Strobelt, H., Luber, J. M., Ouellette, S. B., Azhir, A., Kumar, N., Hwang, J., Lee, S., Alver, B. H., Pfister, H., Mirny, L. A., Park, P. J., & Gehlenborg, N. (2018). HiGlass: Web-based visual exploration and analysis of genome interaction maps. Genome Biology, 19(1), 125. 10.1186/s13059-018-1486-1

Klein, N. K., & Brown, W. M. (1994). Intraspecific Molecular Phylogeny in the Yellow Warbler (Dendroica petechia), and Implications for Avian Biogeography in the West Indies. Evolution, 48(6), 1914. 10.2307/2410517

Korlach, J., Gedman, G., Kingan, S. B., Chin, C.-S., Howard, J. T., Audet, J.-N., Cantin, L., & Jarvis, E. D. (2017). De novo PacBio long-read and phased avian genome assemblies correct and add to reference genes generated with intermediate and short reads. GigaScience, 6(10), gix085. 10.1093/gigascience/gix085

Krueper, D. J. (1996). Effects of livestock management on Southwestern riparian ecosystems. In: Shaw, Douglas W.; Finch, Deborah M., Tech Coords. Desired Future Conditions for Southwestern Riparian Ecosystems: Bringing Interests and Concerns Together. 1995 Sept. 18-22, 1995; Albuquerque, NM. General Technical Report RM-GTR-272. Fort Collins, CO: U.S. Department of Agriculture, Forest Service, Rocky Mountain Forest and Range Experiment Station. p. 281-301., 272, 281–301.

Lamichhaney, S., Han, F., Berglund, J., Wang, C., Almén, M. S., Webster, M. T., Grant, B. R., Grant, P. R., & Andersson, L. (2016). A beak size locus in Darwin’s finches facilitated character displacement during a drought. Science, 352(6284), 470–474. 10.1126/science.aad8786

Li, H. (2013). Aligning sequence reads, clone sequences and assembly contigs with BWA-MEM (arXiv:1303.3997). arXiv. 10.48550/arXiv.1303.3997

Machkour-M’Rabet, S., Santamaría-Rivero, W., Dzib-Chay, A., Cristiani, L. T., & MacKinnon-Haskins, B. (2023). Multi-character approach reveals a new mangrove population of the Yellow Warbler complex, Setophaga petechia, on Cozumel Island, Mexico. PLOS ONE, 18(6), e0287425. 10.1371/journal.pone.0287425

Manni, M., Berkeley, M. R., Seppey, M., Simão, F. A., & Zdobnov, E. M. (2021). BUSCO Update: Novel and Streamlined Workflows along with Broader and Deeper Phylogenetic Coverage for Scoring of Eukaryotic, Prokaryotic, and Viral Genomes. Molecular Biology and Evolution, 38(10), 4647–4654. 10.1093/molbev/msab199

Mérot, C., Oomen, R. A., Tigano, A., & Wellenreuther, M. (2020). A Roadmap for Understanding the Evolutionary Significance of Structural Genomic Variation. Trends in Ecology & Evolution, 35(7), 561–572. 10.1016/j.tree.2020.03.002

Milot, E., Gibbs, H. L., & Hobson, K. A. (2000). Phylogeography and genetic structure of northern populations of the yellow warbler (Dendroica petechia). Molecular Ecology, 9(6), 667–681. 10.1046/j.1365-294x.2000.00897.x

Peona, V., Blom, M. P. K., Xu, L., Burri, R., Sullivan, S., Bunikis, I., Liachko, I., Haryoko, T., Jønsson, K. A., Zhou, Q., Irestedt, M., & Suh, A. (2021). Identifying the causes and consequences of assembly gaps using a multiplatform genome assembly of a bird-of-paradise. Molecular Ecology Resources, 21(1), 263–286. 10.1111/1755-0998.13252

Phillips, S. E., Hamilton, L. P., & Kelly, P. A. (2005). Assessment of habitat conditions for the Riparian Brush Rabbit on the San Joaquin River National Wildlife Refuge, California.

Poff, B., Koestner, K. A., Neary, D. G., & Merritt, D. (2012). *Threats to western United States riparian ecosystems: A bibliography* (RMRS-GTR-269). U.S. Department of Agriculture, Forest Service, Rocky Mountain Research Station. 10.2737/RMRS-GTR-269

Prasad, A., Lorenzen, E. D., & Westbury, M. V. (2022). Evaluating the role of reference-genome phylogenetic distance on evolutionary inference. Molecular Ecology Resources, 22(1), 45–55. 10.1111/1755-0998.13457

Ramírez, F., Bhardwaj, V., Arrigoni, L., Lam, K. C., Grüning, B. A., Villaveces, J., Habermann, B., Akhtar, A., & Manke, T. (2018). High-resolution TADs reveal DNA sequences underlying genome organization in flies. Nature Communications, 9(1), Article 1. 10.1038/s41467-017-02525-w

Ranallo-Benavidez, T. R., Jaron, K. S., & Schatz, M. C. (2020). GenomeScope 2.0 and Smudgeplot for reference-free profiling of polyploid genomes. Nature Communications, 11(1), Article 1. 10.1038/s41467-020-14998-3

Rhie, A., McCarthy, S. A., Fedrigo, O., Damas, J., Formenti, G., Koren, S., Uliano-Silva, M., Chow, W., Fungtammasan, A., Gedman, G. L., Cantin, L. J., Thibaud-Nissen, F., Haggerty, L., Lee, C., Ko, B. J., Kim, J., Bista, I., Smith, M., Haase, B., … Jarvis, E. D. (2020). Towards complete and error-free genome assemblies of all vertebrate species. bioRxiv, 2020.05.22.110833. 10.1101/2020.05.22.110833

Rhie, A., McCarthy, S. A., Fedrigo, O., Damas, J., Formenti, G., Koren, S., Uliano-Silva, M., Chow, W., Fungtammasan, A., Kim, J., Lee, C., Ko, B. J., Chaisson, M., Gedman, G. L., Cantin, L. J., Thibaud-Nissen, F., Haggerty, L., Bista, I., Smith, M., … Jarvis, E. D. (2021). Towards complete and error-free genome assemblies of all vertebrate species. Nature, 592(7856), Article 7856. 10.1038/s41586-021-03451-0

Salgado-Ortiz, J., Marra, P. P., Sillett, T. S., & Robertson, R. J. (2008). Breeding Ecology of the Mangrove Warbler (Dendroica Petechia Bryanti) and Comparative Life History of the Yellow Warbler Subspecies ComplexEcología Reproductiva de Dendroica petechia bryanti y Comparación de los Rasgos de Historia de Vida de las Subespecies del Complejo de Dendroica petechiaSalgado-Ortiz et al.Breeding Ecology of the Mangrove Warbler. The Auk, 125(2), 402–410. 10.1525/auk.2008.07012

Sauer, J. R., Hines, J. E., Fallon, J. E., & Pardiek, K. L. (2014). *The North American Breeding Bird Survey* (Version 02.19) [dataset]. U.S. Geological Survey Patuxent Wildlife Research Center.

Shaffer, H. B., Toffelmier, E., Corbett-Detig, R. B., Escalona, M., Erickson, B., Fiedler, P., Gold, M., Harrigan, R. J., Hodges, S., Luckau, T. K., Miller, C., Oliveira, D. R., Shaffer, K. E., Shapiro, B., Sork, V. L., & Wang, I. J. (2022). Landscape Genomics to Enable Conservation Actions: The California Conservation Genomics Project. *Journal of Heredity*, esac020. 10.1093/jhered/esac020

Shuford, W. D., Gardali, T., Western Field Ornithologists, California, & Department of Fish and Game. (2008). California bird species of special concern: A ranked assessment of species, subspecies, and distinct populations of birds of immediate conservation concern in California. Western Field Ornithologists ; California Dept. of Fish and Game. http://books.google.com/books?id=9INFAQAAIAAJ

Sim, S. B., Corpuz, R. L., Simmonds, T. J., & Geib, S. M. (2022). HiFiAdapterFilt, a memory efficient read processing pipeline, prevents occurrence of adapter sequence in PacBio HiFi reads and their negative impacts on genome assembly. BMC Genomics, 23(1), 157. 10.1186/s12864-022-08375-1

Toews, D. P. L., Taylor, S. A., Vallender, R., Brelsford, A., Butcher, B. G., Messer, P. W., & Lovette, I. J. (2016). Plumage Genes and Little Else Distinguish the Genomes of Hybridizing Warblers. Current Biology, 26(17), 2313–2318. 10.1016/j.cub.2016.06.034

Uliano-Silva, M., Nunes, J. G. F., Krasheninnikova, K., & McCarthy, S. A. (2021). *marcelauliano/MitoHiFi: Mitohifi_v2.0* [Computer software]. Zenodo. 10.5281/zenodo.5205678

Wellenreuther, M., & Bernatchez, L. (2018). Eco-Evolutionary Genomics of Chromosomal Inversions. Trends in Ecology & Evolution, 33(6), 427–440. 10.1016/j.tree.2018.04.002

Wilson, C. M., & Holberton, R. L. (2004). Individual Risk Versus Immediate Reproductive Success: A Basis for Latitudinal Differences in the Adrenocortical Response to Stress in Yellow Warblers (Dendroica Petechia). The Auk, 121(4), 1238–1249. 10.1093/auk/121.4.1238

