## Supplemental Figure S1 for "A highly contiguous genome assembly for the Yellow Warbler (*Setophaga petechia*)"

This PDF file includes:

Figure S1

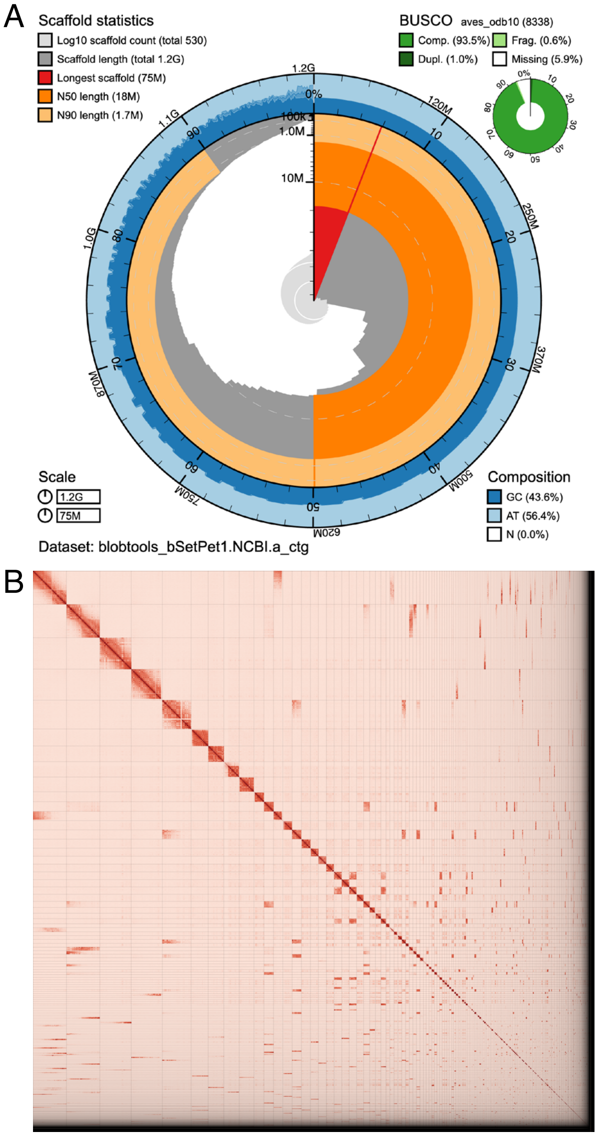


**Figure S1.** Visual overview of genome assembly metrics for alternate assembly (bSetPet1.0.a). A) BlobToolKit Snail plot showing a graphical representation of the quality metrics presented in Table 2. The plot circle represents the full size of the assembly. From the inside-out, the central plot covers scaffold and length-related metrics. The central light gray spiral shows the cumulative scaffold count with a white line at each order of magnitude. The red line represents the size of the longest scaffold; all other scaffolds are arranged in size-order moving clockwise around the plot and drawn in gray starting from the outside of the central plot. Dark and light orange arcs show the scaffold N50 and scaffold N90 values. The outer light and dark blue ring show the mean, maximum, and minimum GC versus AT content at 0.1% intervals (Challis et al. 2020). B) Omni-C contact map for the alternate genome assembly generated with PretextSnapshot. Omni-C contact maps translate proximity of genomic regions in 3D space to contiguous linear organization. Each cell in the contact map corresponds to sequencing data supporting the linkage (or join) between 2 such regions. Scaffolds are separated by black lines and higher density corresponds to higher levels of fragmentation.
